# Integrative model to coordinate the oligomerization and aggregation mechanisms of CCL5

**DOI:** 10.1101/755322

**Authors:** Yi-Chen Chen, Siou-Pei Chen, Jin-Ye Li, Pei-Chun Chen, Yi-Zong Lee, Kun-Mou Li, Raz Zarivach, Yuh-Ju Sun, Shih-Che Sue

## Abstract

The CC-type chemokine ligand 5 (CCL5) is involved in the pathogenesis of many inflammatory conditions. The oligomerization and aggregation of CCL5 are considered to be responsible for its inflammatory properties. The CC-type dimer acts as the basic unit to constitute the oligomer. However, the structural basis of CCL5 oligomerization remains controversial. In this study, NMR and biophysical analyses proposed evidence that no single dimer-dimer interaction dominates in the oligomerization process of CCL5. CCL5 could oligomerize alternatively through two different interactions, E66-K25 and E66-R44/K45. In addition, a newly determined trimer structure reported an interfacial interaction through the N-terminal ^12^FAY^14^ sequence. The interaction contributes to aggregation and precipitation. In accordance with the observations, an integrative model explains the CCL5 oligomerization and aggregation process. CCL5 assembly consists of two types of dimer-dimer interactions and one aggregation mechanism. For full-length CCL5, the molecular accumulation triggers oligomerization through the E66-K25 interaction, and the ^12^FAY^14^ interaction acts as a secondary effect to derive aggregation. The E66-R44/K45 interaction dominates in CCL5 N-terminal truncations. The interaction would lead to filament-like formation in solution.

## Introduction

The CC-type chemokine CCL5 (also known as RANTES) is an 8 kDa inflammatory factor that contains 68 residues. CCL5 is considered to be released from activated immune cells, such as monocytes, macrophages and T lymphocytes (Mariani, Pulsatelli et al., 2002). More recent evidence has shown that CCL5 can also be secreted from activated platelets, epithelial cells and dermal fibroblasts (Niwa, Akamatsu et al., 2001, Schröder, 1995, Terada, Maesako et al., 1997). The chemotaxis activity of CCL5 makes it a very substantial chemoattractant for immune cells. In addition, CCL5 interacts with many inflammatory cytokines, further extending and exacerbating the inflammation reaction. Therefore, CCL5 is associated with a wide range of immune-mediated diseases, including transplant rejection, ischemia-reperfusion injury, atherosclerosis, rheumatoid arthritis, atopic dermatitis, asthma, airway inflammatory disorders, HIV-1 infection and cancers (Arnaud, Beguin et al., 2011, Gerard & Rollins, 2001, Niwa et al., 2001, Roscic-Mrkic, Fischer et al., 2003, Udi, Schuler et al., 2013). The broad effect of CCL5 is correlated with the binding diversity in recognizing several G protein-coupled receptors (GPCRs), including CCR1, CCR3, CCR4 and CCR5 (Appay & Rowland-Jones, 2001, Pakianathan, Kuta et al., 1997, Soria & Ben-Baruch, 2008, Udi et al., 2013).

CCL5 also interacts strongly with glycosaminoglycans (GAGs) that are distributed in the extracellular matrix and on the endothelial cell surface (Kuschert, Coulin et al., 1999, Martin, Blanpain et al., 2001, Singh, Kett et al., 2015). A common feature of GAGs is the large population of sulfate and carboxylate groups, which makes GAGs highly acidic and provides an electrostatic interaction with cationic proteins, including chemokines. CCL5 has a high affinity for heparin and heparan sulfate (Shaw, Johnson et al., 2004, Singh et al., 2015). The binding has been reported to be mediated through a basic heparin-binding motif, where a ^44^RKNR^47^ sequence located in the 40s loop acts as a primary heparin-binding region and a ^55^KKWVR^59^ sequence located in the 50s loop acts as a secondary binding region (Martin et al., 2001, Proudfoot, Fritchley et al., 2001, Segerer, Johnson et al., 2009).

The general concept is that chemokines bind to GAGs through high-order oligomer formation while chemokine monomers bind to protein receptors (Appay, Brown et al., 1999, Bacon, Premack et al., 1995, Chen, Wu et al., 2017, Wang, Sharp et al., 2013). CCL5 has also been reported to act via two signal transduction pathways in T cells. At low concentrations, CCL5 activates GPCRs to regulate transient calcium influx and chemotaxis. At higher concentrations, CCL5 induces a GPCR-independent pathway. The concentration-dependent effect explains the functional diversities of CCL5 (Appay et al., 1999, Bacon, Szabo et al., 1996, Szabo, Butcher et al., 1997). A remarkable case revealed that native CCL5 inhibited HIV infection but became an enhancer of HIV-1 infection when the concentration was high (Cocchi, DeVico et al., 1995, Czaplewski, McKeating et al., 1999, Gordon, Muesing et al., 1999). Modified variants of CCL5 such as AOP-CCL5 and 5P12-CCL5 with less oligomerization ability could be used in inhibiting HIV infection (Wang, Sieg et al., 2013, Wiktor, Hartley et al., 2013, Wilken, Hoover et al., 1999). These analogs compete with HIV in binding CCR5 through monomeric formations and have no effect on forming high-order oligomers. The phenomena support the relevance of studying both oligomeric and monomeric states.

CCL5 has a strong tendency to form noncovalent self-aggregation near neutral pH. Because of the strong aggregation property, there is less structural information for CCL5 under physiological conditions. Most studies were performed under acidic conditions because the aggregates would be dissociated into low-order oligomers and even dimers and monomers (Duma, Haussinger et al., 2007, Hoover, 2000, Skelton, Aspiras et al., 1995). There are two published CCL5 oligomer structures. A tetramer model (PDB code 2L9H) was derived under weak acidic conditions through a hybrid method involving NMR, SAXS and molecular simulation (Wang, Watson et al., 2011). The structural element β1 makes cross-unit contacts with the C-terminus through electrostatic and hydrophobic interactions. Residue K25 makes an intermolecular salt bridge with E66 and two aromatic residues, Y27 and F28, provide hydrophobic contacts with L65 and I62. The interaction can be propagated resulting in an elongated polymeric model among dimer units. Another hexamer structure (PDB code 5CMD) determined by X-ray crystallography was observed using an N-terminal truncation, CCL5(4-68) (Liang, Triandafillou et al., 2016). The N-terminal truncation represents a CCL5 analog produced by N-terminal proteolytic processing (Lim, Burns et al., 2005, Lim, Lu et al., 2006). The structure has been extended to a rod-shaped, double-helical polymer. Residue E26 forms a salt bridge with R47, whereas E66 is located at the positive pocket surrounding R44 and K45. However, the polymer structure shows less probability of the 40s loop to interact with heparin.

The two structures established by different strategies exhibit no similarity. Neither can comprehensively explain all observed structural and functional correlations. The mechanism of how CCL5 forms high-order oligomers is still unclear. In this study, we attempt to answer this question after integrating the following observations. First, after determining the mouse CCL5 dimer structure, the dimer unit was considered the unit for constituting mammalian CCL5 oligomers. Second, by monitoring the different content of aggregation and precipitation upon increasing the pH, we clarified three phases in the CCL5 aggregation process: monomer/dimer, soluble oligomer/polymer and insoluble aggregation/precipitation. Third, we developed a strategy to trap the CCL5 oligomer and prepared a CCL5 trimer (Chen, Li et al., 2018). The solved structure reports a new interface in modulating CCL5 aggregation and precipitation. Finally, we examined the residues currently known to control CCL5 oligomerization to validate the three oligomeric structures, including the current trimer structure. We built an integrative model to satisfy all current observations and explain the oligomerization and aggregation mechanisms of CCL5.

## Results

### Mammalian CCL5s

Human CCL5 (hCCL5) is the only CCL5 with a 3D structure (Chung, Cooke et al., 1995, Hoover, 2000, Shaw et al., 2004). Although CCL5s in other organisms with high sequence identity with hCCL5, there is no structural information about them. It is still worth studying the other CCL5s and comparing the structures to the hCCL5 structure. The mouse model is generally used for *in vivo* and preclinical tests. Thus, we studied mouse CCL5 (mCCL5) (Figure 1). In hCCL5, dimer formation has been considered the structural unit, and the dimer structure becomes the basis for constituting the high-order oligomer structure. Acidic conditions dissociate hCCL5 oligomeric formation and preserve dimer formation. We tested mCCL5 and found that mCCL5 exhibits the same property. The dimer configuration was confirmed by NMR ^1^H-^15^N HSQC spectra (Figure 1B and Expanded view Figure EV1). We solved the dimer structure using the programs CYANA3.0 (Guntert, 2004) and X-PLOR (Schwieters, Kuszewski et al., 2003). The dimer structure adopts a typical CC-type dimer formation. Twenty low-energy dimer structures were selected from 100 calculated structures and subsequently performed energy minimization. The details of structural determination are summarized in Appendix and Table EV1. The energy and structure statistics are listed in Table EV2. The root-mean-square deviation (RMSD) values of residues 6-66 were 0.70 ± 0.24 Å for the backbone atoms and 1.01 ± 0.21 Å for the heavy atoms. The Ramachandran plot showed no residues distributed in disallowed regions (Figure EV2). The main structural features are an unstructured N-terminus (residues 1-6), a short anti β-strand that forms an interfacial region with another unit (residues 9-11), a single turn of 3_10_ helix (residues 22-24), a three-antiparallel β-sheet (β1, residues 26-30; β2, residues 40-44; β3, residues 47-52) and a C-terminal α-helix (α1, residues 57-66) (Figure 1C). The α-helix packs onto the β-sheet surface to stabilize the anti-parallel β-sheet. The structure contains two disulfide bonds, Cys 10 with Cys 34 and Cys 11 with Cys 50.

**Figure 1.**
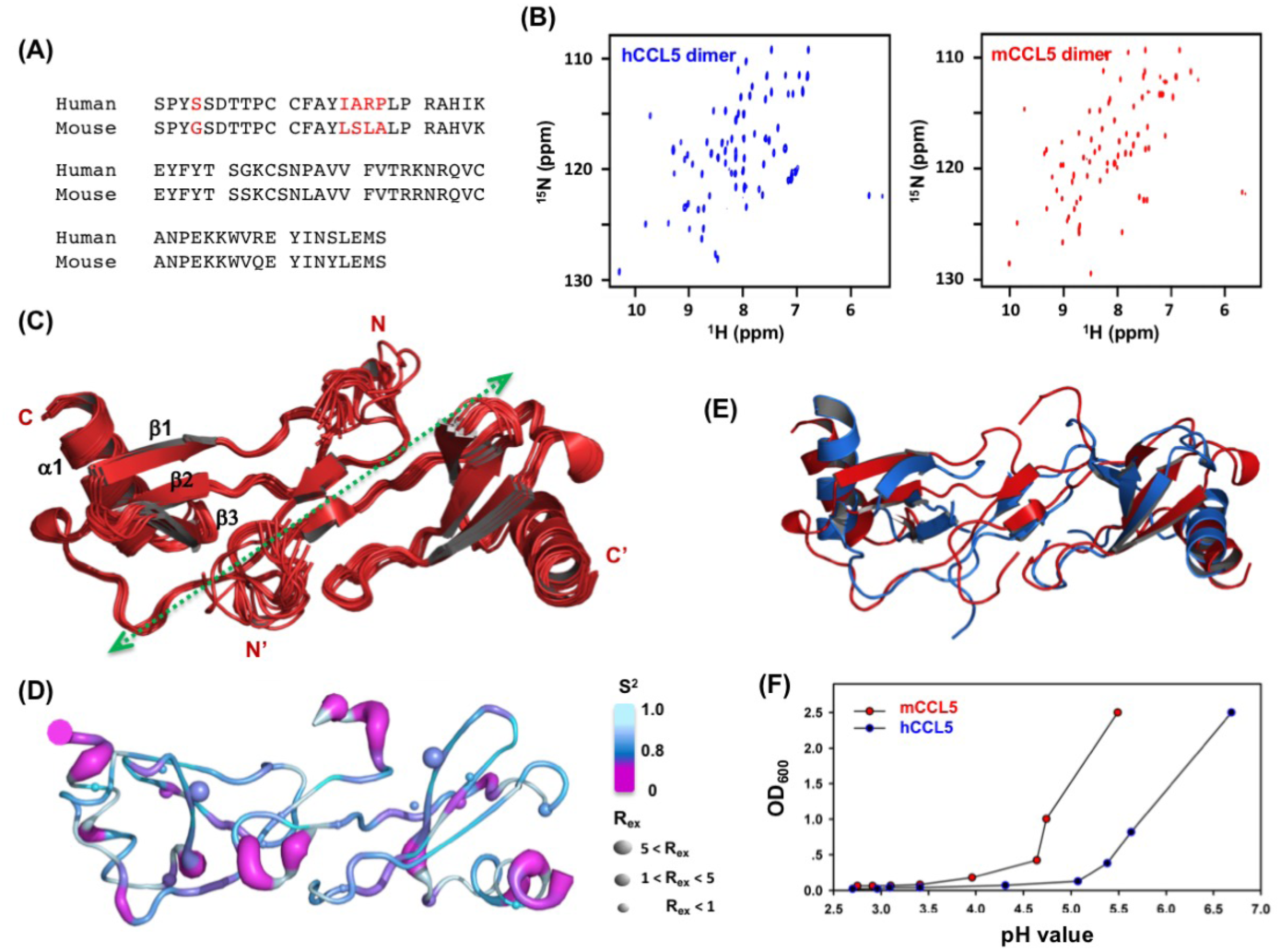
Solution structure of mouse CCL5. (A) Sequence alignment between human CCL5 (hCCL5) and mouse CCL5 (mCCL5). (B) Comparison of the ^1^H,^15^N-HSQC spectra of hCCL5 (blue) and mCCL5 (red); 0.2 mM proteins were in 25 mM sodium acetate, pH 4.0 (hCCL5) or pH 3.2 (mCCL5), 150 mM NaCl and 10% D_2_O at 298K. (C) Superimposed NMR structures of mCCL5. Twenty lowest energy structures were superimposed by the best-fit superposition of the backbone atoms. (D) Model-free analysis based on ^15^N relaxation parameters. Structural mapping of the dynamic parameters of S^2^ and R_ex_ by a sausage model. Residues with high order parameters (S^2^ > 0.9) indicate structural rigidity, whereas residues with R_ex_ perform µs to ms time-scale conformational exchange. (E) Structure comparison between hCCL5 (cyan, PDB code 1EQT, 1.6 Å resolution) and mCCL5 (red). (F) The pH-dependent turbidity assay of hCCL5 and mCCL5; 0.2 mM proteins were in 25 mM sodium acetate, 150 mM NaCl at 298 K.

The NMR ^15^N relaxation data, including R_1_, R_2_, and heteronuclear NOEs, report the backbone dynamics (Figure EV3). The mean relaxation parameters were R_1_ = 1.27 s^-1^ and R_2_ = 11.31 s^-1,^ and the R_2_/R_1_ parameter reported an overall rotational correlation time of 9.08 ns, corresponding to a dimer. By using FastModelFree fitting, the backbone dynamics parameters S^2^, τe and R_ex_ were derived, representing the order parameter, internal motion and chemical exchange, respectively (Figure EV3). We mapped the order parameter, S^2^ and R_ex_, on the mCCL5 structure (Figure 1D). The model exhibits structural rigidity, with most S^2^ values higher than 0.9. The N-terminus, the end of α1 and the 50s loop (residue 51-54) have regional flexibility. A total of 10 residues exhibit R_ex_, which are distributed sparsely on the structure (Figure 1D).

### Comparison between mouse and human CCL5

The two CCL5s share similar dimeric properties that both adopt typical CC-type dimer formations, and the N-terminal interfacial region is responsible for dimerization (Hoover, 2000). The averaged mCCL5 structure was superimposed on the hCCL5 crystal structure (Figure 1E). The mCCL5 and hCCL5 monomer structures had small RMSD values of 1.044 Å. The corresponding RMSD value between the two dimers was increased to 1.973 Å. The difference mainly comes from the orientation between the two monomers, indicating structural flexibility in the dimer interface. In acidic conditions, dimer formation is dominant with stability. Combining the cases of hCCL5 and mCCL5, we expect the dimer property to be extended in the other mammalian CCL5s.

### Precipitation and aggregation property

We next examined the aggregation behavior of hCCL5 and mCCL5 under different pH conditions associated with the turbidity assay. Turbidity is reported by light scattering, reflected in visible absorption. The experiments reveal the molecular precipitation tendency. Both CCL5s showed pH-dependent turbidity in which no huge complex was detected under acidic conditions (pH < 3.5), and turbidities were dramatically increased when the pH value reached 4.5 for mCCL5 and 5.5 for hCCL5 (Figure 1F). The turbidities continued to increase when the pH was close to a neutral condition, and the maximum absorbances (OD_600_) were > 2.5. The results suggest similar precipitation properties in the two chemokines. However, the isoelectric points of mCCL5 and hCCL5 were 8.76 and 9.24, respectively. Thus, the phenomenon is not due to the isoelectric precipitation but instead, based on the property of protein assembly. Notably, mCCL5 showed higher sensitivity upon the pH increase. The difference was derived from the sequence difference between the two CCL5s, such as positions 4, 15-18, 24, 32, 37, 45 and 58 (Figure 1A). The major sequence discrepancy occurs in the N-terminal region. To examine the correlation, we substituted the mCCL5 sequence with the hCCL5 sequence at positions 4 and 15-18. The derived mutants, including mCCL5-^4^S, mCCL5-^17^RP^18^ and mCCL5-^15^IARP^18^, had increased precipitation propensity at neutral pH (Figure EV4), behaving more similar to hCCL5. The precipitation of CCL5 was modulated by the N-terminal portion of CCL5.

### Phases of CCL5 oligomerization

Aggregation and precipitation are considered consequences of massive oligomerization. We compared the turbidity assay and NMR ^1^H-^15^N HSQC spectra (Figure 2). Taking hCCL5 as an examined case, three phases were clarified. In phase I, when the pH was lower than pH 4.0, no turbidity was detected, and NMR revealed the dimer configuration together with trace amounts of monomer. When the pH was increased to 4.0-5.0, the turbidity remained low, and no precipitation was observed. However, the signal intensities in HSQC were dramatically attenuated, indicating severe oligomerization. In the phase II, the oligomerized complex had good solubility, and the solution was still transparent. When the pH was higher than 6.0, the turbidity rapidly increased, and almost no signal was detected in HSQC. The CCL5 molecules were not only polymerized but also started to aggregate and formed an insoluble complex. We conclude that insoluble aggregation/precipitation occurred in the third phase.

**Figure 2.**
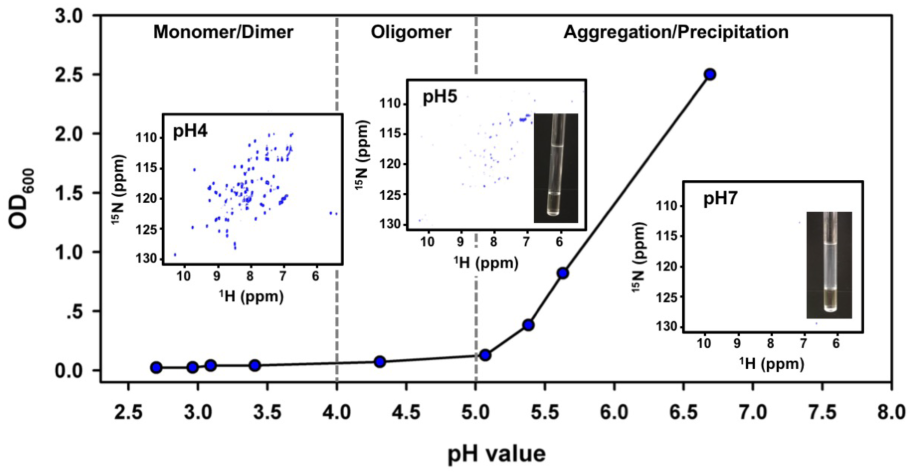
Three phases of CCL5 aggregation, classified from the pH-dependent turbidity assay and ^1^H,^15^N-NMR HSQC spectra.

### CCL5 trimer structure

We intend to determine the CCL5 oligomer structure under neutral conditions. The CCL5-E66S mutant has been known to adopt stable dimer formation in a wide pH range (Czaplewski et al., 1999, Wang et al., 2011). Mixing the E66S mutant with native CCL5 reduced the propensity of CCL5 oligomerization and aggregation. The NMR HSQC experiment revealed that the addition of hCCL5-E66S caused the broadened resonances of ^15^N-hCCL5 to become intense and dispersive, indicating dissociation from the hCCL5 complex (Figure 3). Meanwhile, in the reverse titration, the resonances of ^15^N-hCCL5-E66S were of attenuated intensity, indicating an association with the hCCL5 complex to behave like a “big” molecule. Therefore, a strategy of preparing CCL5 oligomers by mixing native hCCL5 with the E66S mutant was developed (Figure 3). Under a 1:2-1:3 molar ratio, we trapped a CCL5 trimer. The size distribution was confirmed by size-exclusion chromatography (SEC) and SAXS experiments (Chen et al., 2018). The trimer sample was crystallized, and the structure was determined. The data collection and refinement statistics are listed in Table EV3.

**Figure 3.**
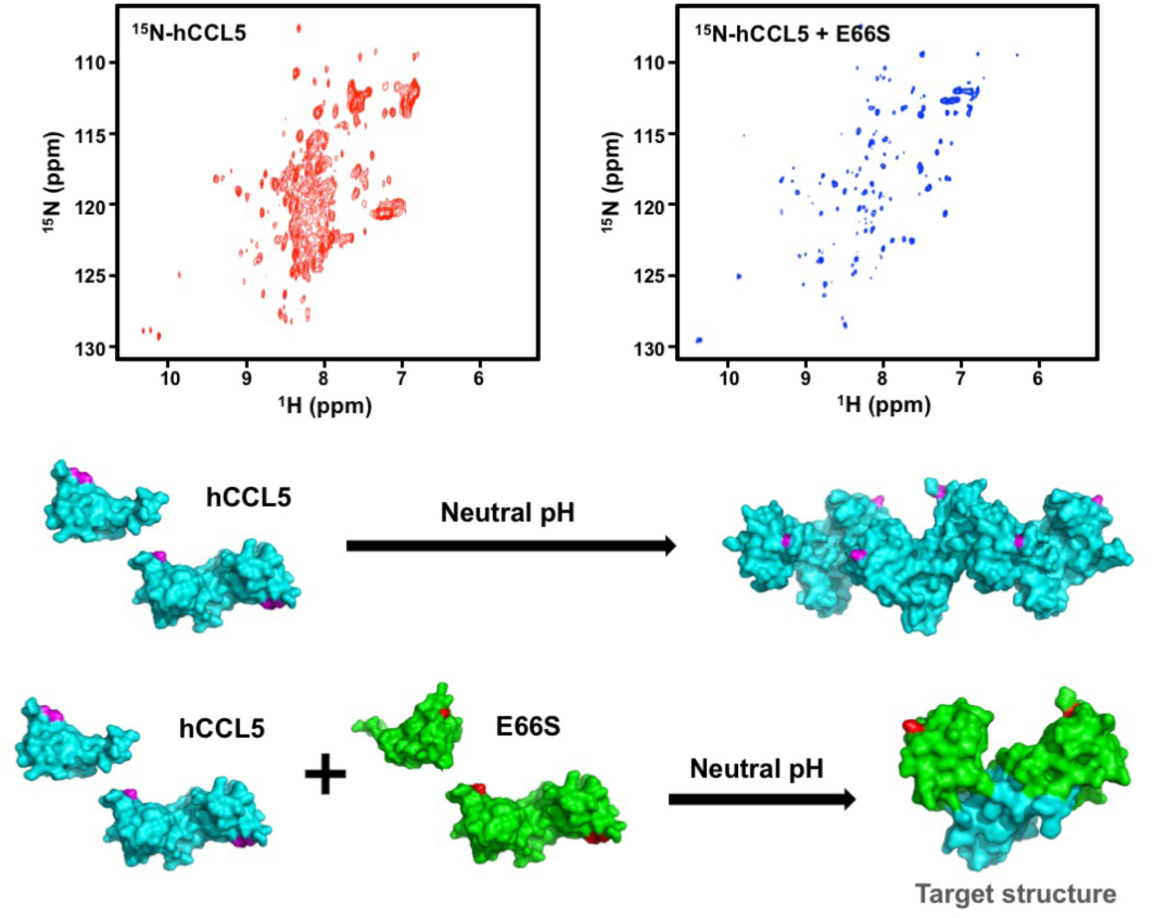
The strategy to prepare CCL5 oligomers. The CCL5-E66S mutant was mixed with native CCL5 to reduce the oligomerization propensity. NMR ^1^H,^15^N-HSQC spectra of ^15^N-hCCL5 without or with hCCL5-E66S (molar ratio of 1:3) in 25 mM sodium acetate, pH 5. The two proteins were mixed in an acid condition, followed by exchanging to pH 5.

The crystal structure of hCCL5 is a heterotrimer in an asymmetric unit, which contains three subunits indicated by monomers A, B and C in Figure 4A. Monomers A and C are hCCL5-E66S, and monomer B is native CCL5. The three monomeric subunits have nearly identical structures, and the electron densities of the residues are generally satisfactory except for residues 1-5 and 70 in A, 1-2 and 67-70 in B, and 1-2 and 70 in C. The main features of the secondary structural elements are the same as the previously determined structure (PDB code 1EQT) and the current-determined mCCL5 structure. In the structure, monomers B and C form a typical CC type dimer, and A, acting as a peripheral subunit, exhibits an atypical contact with monomer B (Figure 4A). The atypical interface contains interactions mainly through Y14, where short anti-parallel β-sheet forms from the ^12^FAY^14^ sequences of monomers A and B. Intermolecular backbone hydrogen bonds occur between the residues F12 to A16’ and Y14 to Y14’. Extra hydrophobic interactions occur between A13 to Y14’, I15 to A16’ (Figure EV5).

**Figure 4.**
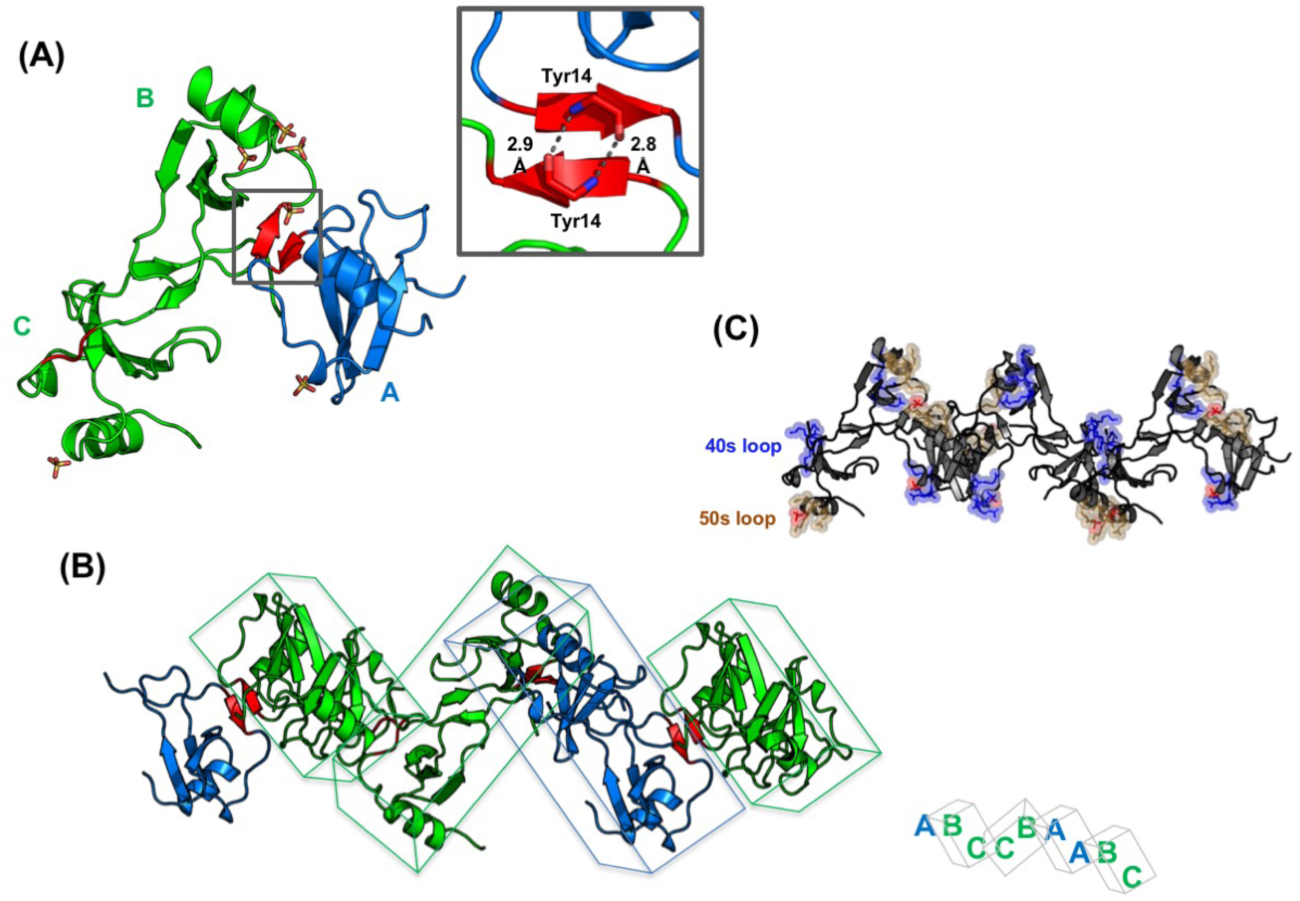
Structure of the hCCL5 trimer. (A) The hCCL5 trimer contains a CC-type dimer and a peripheral binding monomer. The ^12^FAY^14^ interface between the monomer (chain A) and dimer (chains B and C) is colored red, and the detailed interactions are shown in the box. Intermolecular hydrogen bonds through Y14 are represented by dashed lines. The crystallized sulfate ions are indicated by sticks. (B) A W-shaped oligomer model in which CCL5 is alternatively polymerized through the CC-type dimer interface and the ^12^FAY^14^ interface. (C) The relative distribution of the 40s and 50s loops in the model. The sulfate ions, indicated by red, represent the potential sites for heparin binding.

The identified interface allows for the creation of a new model for the CCL5 oligomer. The typical CC-type dimer (units B and C) can link with another dimer through the new interface, constituted by the motif ^12^FAY^14^. In the model, the N-terminal residues of P2 and S4 additionally extend to contact the hydrophobic residues of I15 and A13 of another dimeric unit. The additional interactions might stabilize the polymer structure. By repeating the interaction, a W-shaped oligomer model was generated (Figure 4B). There were six sulfate ions identified in one trimer structure, located near the 40s and 50s loops (Figure 4A). The sulfate ions closely interact with neighboring positive residues and occupy the potent heparin-binding sites. One sulfate ion located near the 40s loop interacts with R44, K45 and R47 in unit A, and the remaining sulfate ions near the 50s loop interact with K55 and K56 and partially interact with R59 in units A, B and C. The modeled polymer fully exposes the 40s and 50s loops, benefitting GAG binding (Figure 4C).

### Structural motif ^12^FAY^14^

The new interfacial interaction involved the structural motif ^12^FAY^14^. The two aromatic residues are strictly conserved in all mammalian CCL5s (Figure 5A). To examine the relevance, we prepared a mutant, hCCL5-^12^AAA^14^, in which the two aromatic residues are both substituted by Ala. The mutant showed significantly reduced turbidity (Figure 5B). Turbidity only increased when the pH reached the isoelectric point. NMR HSQC spectra demonstrated dispersive resonances in both acidic and neutral conditions (Figure 5C). However, slight line broadening was still observed in the HSQC at pH 7, indicating partial oligomerization. We demonstrated the involvement of ^12^FAY^14^ in CCL5 oligomerization, aggregation and precipitation. The result corresponds to the previous observation of the CCL5 N-terminal portion modulating CCL5 precipitation (Figure EV4).

**Figure 5.**
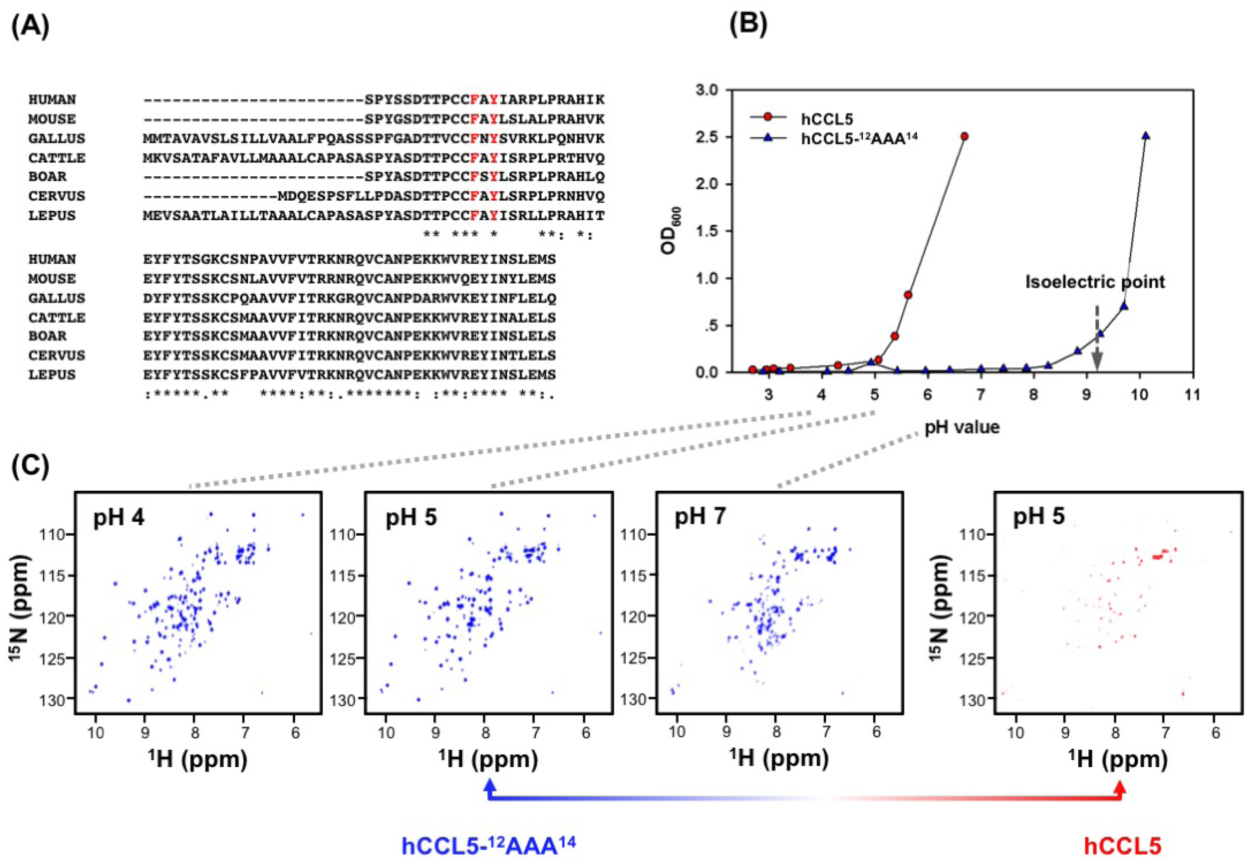
Solution properties of the hCCL5-^12^AAA^14^ mutant. (A) Conserved ^12^FAY^14^ motif in different mammalian CCL5s. (B) The pH-dependent turbidity assay. (C) The corresponding ^1^H,^15^N-HSQC spectra at pH 4, 5 and 7 in 25 mM sodium acetate and 150 mM NaCl. The solution property was compared with native hCCL5.

To precisely define the role of the ^12^FAY^14^ motif, we prepared 4 mutations in which the two aromatic residues were replaced by Ala (with reduced sidechain effect) or Pro (without backbone NH). The substitutions can differentiate the effect from the sidechain hydrophobic interactions or backbone hydrogen bonding. The turbidity assays of the mutants F12A, F12P and Y14A were similar to hCCL5-^12^AAA^14^, in which the precipitation propensities were eliminated (Figure 6). In the Y14P mutant, the precipitation propensity was significantly reduced compared with the native protein but higher than the other mutations. The OD_600_ value of Y14P gently increased when the pH was higher than 6. Since the substations of F12A and Y14A were both effective in reducing precipitation, sidechain hydrophobic interactions are critical. The backbone hydrogen bonding in Y14 contributed a supporting role in the aggregation and precipitation process.

**Figure 6.**
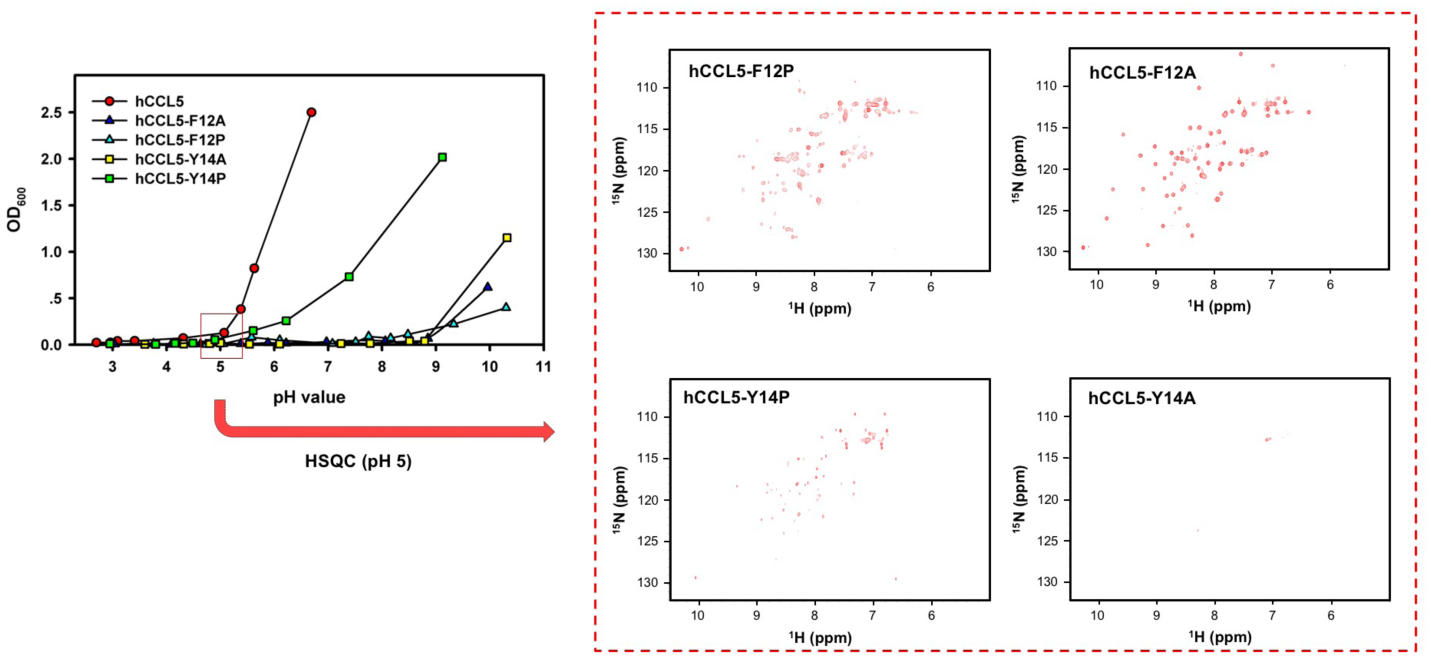
The pH-dependent turbidity assay of hCCL5 single mutations at positions 12 and 14. The corresponding ^1^H,^15^N-HSQC spectra at pH 5.0 were compared to show the different oligomerization properties.

We next performed NMR analysis for these mutants at pH 5.0, at which the soluble CCL5 oligomers readily enter the aggregation/precipitation phase (from phase II to phase III). Compared with the HSQC spectrum of native CCL5, shown in Figure 5C, the mutants F12A and F12P had intense resonances, corresponding to the CCL5 monomer in F12A and monomer/dimer mixture in F12P (Figure 6). The mutations at position 12 caused dimer dissociation and promoted monomer formation, therefore reducing oligomerization. Subsequently, the aggregation phase was prohibited. The mutants Y14A and Y14P demonstrated almost undetectable signals in the HSQC spectra, indicating the formation of a large protein complex. The mutants instead maintained the oligomerization property but exhibited impaired aggregation, as reflected in the low turbidity. Residue Y14 contributed to CCL5 aggregation, not oligomerization.

### The self-associations of different oligomers

Using the same strategy, we further validated other oligomer structures and the currently known residues by using NMR and turbidity assays. The acidic residue, E66, critically controls the CCL5 oligomer. In the determined oligomer structures, the tetramer model (PDB code 2L9H) reported charge-charge interaction between E66 and K25, and the hexamer structure (PDB code 5CMD) revealed the interaction between E66 and two positively charged residues, R44 and K45 (Liang et al., 2016, Wang et al., 2011). We prepared two mutants to evaluate the involvement of these residues.

We first examined the E66-K25 interaction. The mutant hCCL5-E66S induced no turbidity in solution, consistent with the presence of stable dimers in solution (Figure 7A). The mutant hCCL5-K25S also had very low aggregation/precipitation properties (Figure 7A). However, the corresponding NMR HSQC demonstrated extremely weak resonances at pH 6.0, indicating a large molecular size (Figure 7B). The formed complex was too large to be detected by NMR and had a regulated formation that was soluble in solution. This feature was still maintained when the pH reached 7.0, as proven by the transparent NMR sample (Figure 7C). We observed the existence of a soluble polymer with strong self-assembly of the CCL5 dimer. Since the interaction between E66-K25 was abolished, the property most likely represents the behavior derived from the other interactions between E66 and the R44/K45 cluster. This interaction creates polymers with a vast size and regulated formation without precipitation properties. Notably, the N-terminal truncation, hCCL5(5-68), which was reported to adopt a hexamer formation, showed similar behavior to hCCL5-K25S in the two experiments (Figure 7A). According to these features, we suspect that hCCL5-K25S, behaving like hCCL5(5-68), adopts the hexamer model and extends to the rod-shaped polymer in solution.

**Figure 7.**
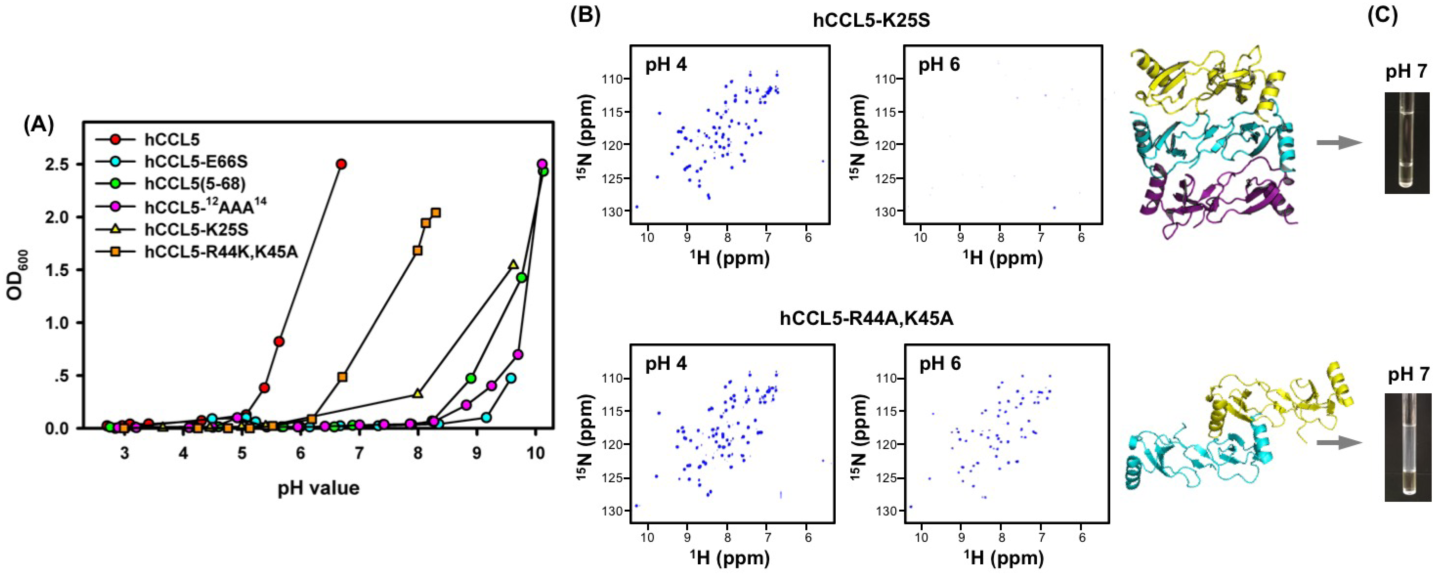
Comparison of pH-dependent turbidity assays and NMR HSQC spectra of different hCCL5 mutants. (A) The pH-dependent turbidity assay. (B) The HSQC spectra at pH 4.0 and 6.0 of hCCL5-K25S, and hCCL5-R44A, K45A and the corresponding oligomer models. (C) The NMR samples of hCCL5-K25S and hCCL5-R44A, K45A at pH 7.0.

Alternatively, the removal of R44 and K45 had less of an effect in reducing CCL5 aggregation, and precipitation still occurred when the pH reached 6.0 (Figure 7A). In contrast to hCCL5-K25S, the corresponding HSQC at pH 6.0 reported a configuration with limited oligomerization, as evidenced by the moderately reduced intensity (Figure 7B). Furthermore, when the pH reached 7.0, the NMR sample showed similar behavior as native CCL5, and massive aggregation was detected in the NMR sample (Figure 7C). Alternatively, the removal of R44 and K45 might reveal features derived from the E66 and K25 interaction. When the oligomerization occurs through the E66-K25 interaction, the initial oligomerization process can be moderate, and the interaction accompanies aggregation and precipitation when the pH approaches neutral. This phenomenon is consistent with the fact that the tetramer model was observed under weakly acidic conditions, corresponding to phase II. Upon increasing the pH, hCCL5 continues to polymerize, but aggregation/precipitation occurs with the process as well.

### TEM analysis

To disclose the different aggregations of CCL5, transmission electron microscopy (TEM) was employed to directly visualize the shape and formation of CCL5 (Figure 8). There are two types of CCL5s identified in cells: full-length CCL5 and its N-terminal truncated variants. We examined these two cases. Full-length CCL5 forms precipitates. At pH 6.0, the hCCL5 complex existed as uniform and homogenous particles. The behavior might be close to phase separation. The images further reported nonuniform precipitation at pH 7.0. No particular formation was identified. However, as observed in the turbidity assay, the N-terminal truncated CCL5 created another type of CCL5 complex. We examined hCCL5(5-68), and the TEM images revealed that the CCL5 analog self-associated into a filament-like formation when the pH reached 6.0-7.0. Again, the image consistently supports the formation of the rod-shaped, double-helical polymer model observed by the corresponding X-ray structure.

**Figure 8.**
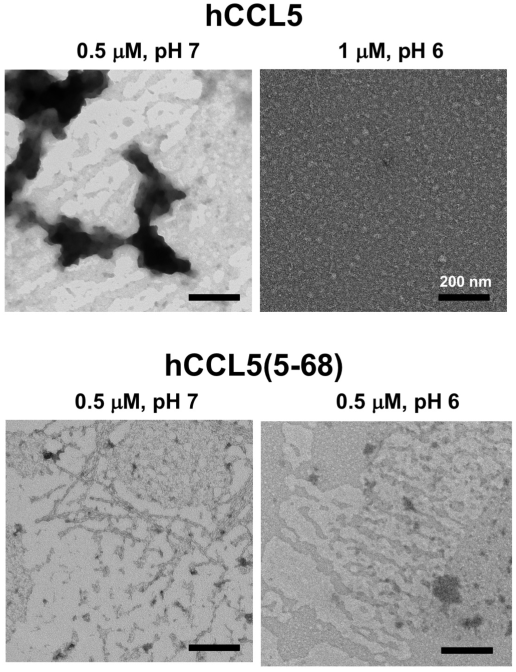
TEM images of full-length hCCL5 and the N-terminal truncation, hCCL5(5-68). The black bar represents 200 nm.

### Heparin binding

Heparin is the most effective GAG that interacts with CCL5. We investigated the heparin-binding of different hCCL5 variants based on the surface plasmon resonance (SPR) method (Figure 9). Biotinylated heparin was immobilized on a C1 chip to eliminate the concern of mass transfer. Different hCCL5s were injected at concentrations of 25-500 nM and flowed over the heparin surface for 300 sec to reach equilibrium. Full-length hCCL5 demonstrated the best binding ability, and the N-terminal truncation, hCCL5(5-68), also contained similar binding affinity. When only the dimerization ability was preserved, weak heparin binding was observed in the hCCL5-E66S mutant. Meanwhile, hCCL5-^12^AAA^14^ showed a similar weak binding ability compared to that of hCCL5-E66S, which corresponded to the deficient oligomerization ability. The binding affinity calculated by steady-state affinity analysis reported a K_D_ value of 415 nM for CCL5, and the oligomer-deficient hCCL5-E66S and hCCL5-^12^AAA^14^ both exhibited a 30-fold reduction in heparin-binding affinity.

**Figure 9.**
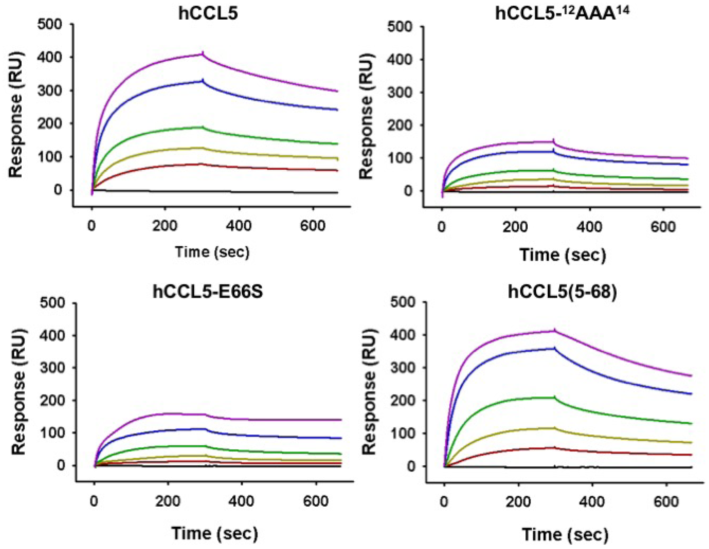
SPR sensorgrams of the binding of hCCL5s to heparin. Sensorgrams of individual CCL5s contain injections of different concentrations (25, 50, 100, 250 and 500 nM) over a heparin-coating chip.

## Discussion

CCL5 shows a high tendency for protein oligomerization and aggregation under physiological conditions. The oligomerized formation has been correlated with cell migration and activation (Appay et al., 1999, Johnson, Kosco-Vilbois et al., 2004, Proudfoot, Handel et al., 2003). Studying the oligomerization mechanism is essential for understanding the chemotaxis activity of CCL5 and how CCL5 binds cell surface receptors. By considering human and mouse CCL5s, the molecules show similarity in the dimeric conformation but have slightly different oligomerization sensitivity toward pH changes. This phenomenon contains a very complicated mechanism, which is reflected in the dependence on protein sequence, pH, ionic strength and protein concentration.

The current study reported a CCL5 trimer structure. The trimer structure revealed the N-terminal conserved ^12^FAY^14^ motif contributing to the interaction between the CCL5 dimer and the other CCL5 monomeric subunit. This trimer structure could be extended to a W-shaped oligomer by alternatively repeating the CC-type dimer interface and the ^12^FAY^14^ interface. This study is not the first case to reveal the ^12^FAY^14^ interface. The ^12^FAY^14^ interface was accidentally reported in another CCL5 variant, P2-CCL5 (Jin, Kagiampakis et al., 2010). P2-CCL5 contains an enhanced ability to inhibit HIV-1 infection. Due to its mutated N-terminal sequence, P2-CCL5 has been characterized as a monomer in solution and demonstrated a diminished capacity to bind GAGs. The P2-CCL5 structure (PDB code 2VXW) surprisingly forms a tetramer in the crystallographic asymmetric unit, in which the ^12^FAY^14^ motif is responsible for dimer-dimer interaction. This structure indirectly validates the potency of the ^12^FAY^14^ site in CCL5 association. Comparing the two structures, the local interactions are almost the same; two intermolecular hydrogen bonds exist between Y14 and Y14’, and the N-terminal end extends to a hydrophobic pocket of another dimeric unit and is stabilized by hydrophobic interactions with residues, such as I15 and A13. However, compared with other determined oligomeric interfaces, the ^12^FAY^14^ interface contains fewer interactions. The interface may not be very stable. In addition, the residue E66, reported to be critical in mediating CCL5 oligomerization, had no contact in this trimer structure. Thus, this trimer formation should not be the dominant model for CCL5 oligomerization. Taken together with the fact that the double mutations F12A and Y14A caused no turbidity or precipitation, we concluded that position 12 contributes to CCL5 dimer association and position 14 is involved aggregation and precipitation. The ^12^FAY^14^ motif plays a role in cooperatively propagating CCL5 oligomers into an aggregated polymeric form. Other interactions are primarily responsible for oligomerization.

As mentioned, there are two determined oligomer structures (Liang et al., 2016, Wang et al., 2011). The charge-charge interactions of E66-K25 and E66-R44/K45 are dominant in stabilizing tetramer model and hexamer structure, respectively. However, none of them can perfectly explain all features of CCL5. We examined the two models and proposed that no single dimer-dimer interaction predominated the oligomerization property. Both interactions could coexist in the process. When the E66-R44/K45 interaction was removed to maintain the E66-K25 interaction in the hCCL5-R44A, K45A mutant, it corresponded to the tetramer model (PDB code 2L9H). The mutant started to oligomerize when the pH increased and still contained similar precipitation properties. This observation complements the fact that the ^12^FAY^14^ motif is fully exposed in the tetramer model to allow recruiting another dimer unit. The conjugation through Y14 has no steric effect against the original dimer-dimer interaction (E66-K25) and might create branched oligomer formation. Therefore, this could be the structural basis for aggregation and precipitation under high concentrations.

In the hCCL5-K25S mutant, the E66-R44/K45 interaction is preserved, and the interaction is important to maintain the hexamer structure (PDB code 5CMD). The behavior of this mutant reflects the feature of hexamer formation. Under neutral conditions, hCCL5-K25S was significantly polymerized, as concluded in the NMR study. Meanwhile, no precipitation was observed. In the hexamer model, the 30s loop stacks on the ^12^FAY^14^ motif, prohibiting further propagation through the interaction of Y14. This result explains the lack of precipitation observed in the hCCL5-K25S mutant. The mutant assemble into the rod-shaped polymer through the E66-R44/K45 interaction. The polymer formation was stable and had good solubility. We noticed that the N-terminal truncated CCL5 demonstrated similar behavior as that of hCCL5-K25S. Moreover, the rod-shaped polymer model was established when studying the N-terminal truncated variant. We suspect that the few flexible N-terminal residues might hinder the stacking of the CCL5 molecules during the crystallization. After removing the residues, the property became dominant and exclusive.

We integrated the information and depicted the full mechanism in Figure 10. The CC-type dimer is the building unit for constituting oligomers and polymers. The monomer-dimer equilibrium occurs when the pH and ionic strength are low (phase I). Upon approaching biological conditions, CCL5 begins to oligomerize through two mechanisms (phase II). In the first pathway, the formed oligomer could comprise a primary dimer-dimer interaction through E66 and K25. The increased concentration of CCL5 starts to create secondary interaction through the ^12^FAY^14^ interface. This additional interaction triggers aggregation and precipitation once the two interactions alternatively conjugate CCL5 molecules (phase III). In the second pathway, the E66-R44/K45 interaction becomes exclusively primary in leading oligomerization. The dimer-dimer interface is stable and allows polymerization into a rod-shaped structure. Without the involvement of the ^12^FAY^14^ site, the structure has no aggregation property.

**Figure 10.**
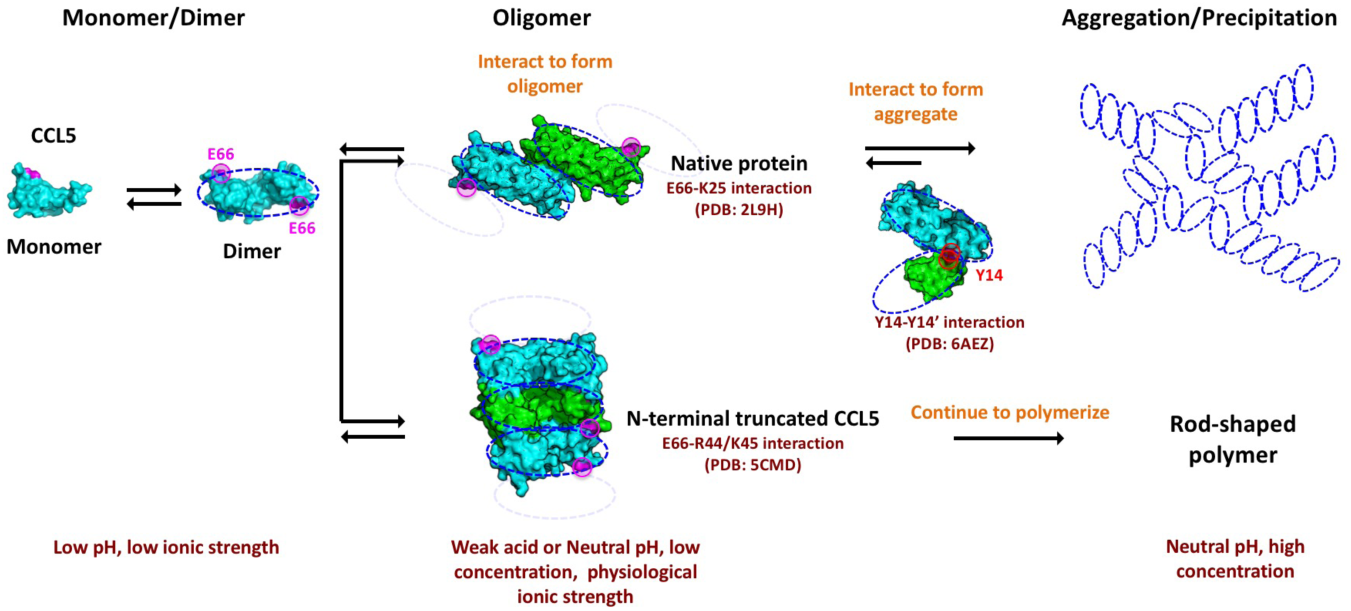
The integrative model illustrates CCL5 oligomerization and aggregation.

The current model brings a very important implication for understanding the functional diversity of active CCL5s. Of the two types of CCL5s identified in cells, full-length CCL5 might favor the first pathway forming oligomers. Severe aggregation would occur when the pH is close to neutral and the concentration is high. SAXS data, reflecting the solution behavior, supported this idea. The structure has good affinity for heparin and generates chemotaxis for immune cells. Alternatively, after proteolytically processing by CD26 or cathepsin G, CCL5 is converted into the N-terminal truncated variants (Lim et al., 2006). The variant adopts the second pathway, forming a rod-shaped oligomer (Liang et al., 2016), which might create a filament-like formation. Although the analogue has heparin-binding affinity, the receptor binding and chemotactic and antiviral activities are generally reduced.

## Materials and Methods

### CCL5 expression and purification

To simplify protein preparation, a methionine residue was incorporated at the CCL5 N-terminus, and the construct was directly expressed in *E. coli*. The DNA sequences of CCL5 from human and mouse were amplified by PCR and ligated into the pET-43.1a(+) vector. When the OD_600_ reached 0.6, the cells were induced by 1 mM IPTG for 5 hours at 37 °C. The cells were harvested by centrifugation (6000 x g, 20 min at 4 °C) and resuspended in lysis buffer (25 mM Tris-HCl pH 8.0, 100 mM NaCl). The mixture was disrupted by a high-pressure homogenizer (GW Technologies, Taiwan). The cell lysate was centrifuged at 15,000 x g for 30 min at 4 °C, which resulted in supernatant and pellet fractions. The pellet was resuspended in denaturing buffer (100 mM Tris, 6 M Gdn-HCl, pH 8.0) and stirred at 25 °C overnight. Cell debris was removed by centrifugation (30,000 x g, 20 min at 4 °C). The supernatant was renatured by 100-fold dialysis into refolding buffer (0.9 M Gdn-HCl, 5 mM cysteine, 5 mM methionine, 100 mM Tris pH 8.0). The solution was mildly stirred at 4 °C (2 liter buffer, twice for 8 hours each). Protein debris was removed by centrifugation (30,000 x g, 20 min at 4 °C). The supernatant containing renatured protein was isolated by HPLC.

### NMR solution structure

To prepare the uniformly single (^15^N) or double isotope (^15^N and ^13^C) labeled protein for NMR analysis, cells were cultured in 10 ml LB medium with 100 µg/ml ampicillin at 37 °C overnight. The cells were collected by centrifugation for 10 min at 5000 x g and resuspended in 1 liter M9 minimum medium (12.8 g Na_2_HPO_4_•7H_2_O, 3 g KH_2_PO_4_, 0.5 g NaCl, 0.1%, 2 mM MgSO_4_, 0.1 mM CaCl_2_, 0.01% thiamine and 80 nM MnCl_2_) with 100 µg/ml ampicillin supplemented with ^15^NH_4_Cl (1 g/L) and [^12^C_6_] D-glucose (2 g/L) or [^13^C_6_] D-glucose (2 g/L) as the sole nitrogen and carbon sources. The expression and purification procedures were the same as described above. To determine the structure, all NMR spectra were acquired on a Bruker AV600 spectrometer at 298 K. The ^15^N and ^15^N,^13^C-labeled samples were prepared in NMR buffer (25 mM sodium acetate pH 3.2, 150 mM NaCl with 10% D_2_O (v/v)) and added to Shigemi NMR tubes (Shigemi Co., LTD). The parameters used in the NMR experiments are listed in Table EV1, and the details of the structure and dynamics determination are described in Appendix.

### X-ray structure determination

The trimer samples were concentrated to 7 mg/ml in 20 mM acetate, pH 5.0 with 150 mM NaCl for initial crystallization screening. Triangle plate-like crystals successfully appeared in the Index HR2-144 screen. The best single-crystal diffraction was obtained at a 1.87 Å resolution on the NSRRC BL13B1 beamline. The crystals had a tetragonal lattice and belonged to space group P4_3_2_1_2, with unit-cell parameters of a=56.6, b=56.6, c=154.1 Å. Matthews coefficient calculations indicated the presence of three molecules per asymmetric unit, matching the observation in the SEC experiment. The phase problem was solved by molecular replacement using the hCCL5 monomer structure (PDB code 1EQT) determined under acidic conditions (pH = 4.6). The molecular replacement indicated that the crystal was in the enantiomorphic space group P4_1_2_1_2. Several rounds of model building and structural refinement were performed to improve the quality. The details of the structural determination are described elsewhere (Chen et al., 2018).

### Turbidity measurement

Proteins were diluted in 25 mM acetate buffer, pH 3.0 with 150 mM NaCl in a final volume of 400 µl, and the protein concentrations were 0.2 mM. The resulting protein solutions were incubated for 1 min at room temperature and ready for analysis. The turbidity derived by protein aggregation was measured by OD_600_ using a UV-visible nanophotometer (Implen) in 1 cm path length optical cells (BL6224, Basic Life). Titrating NaOH into the samples changed the pH from acidic to basic conditions. We monitored the pH values and turbidities for individual measurements.

### Transmission electron microscope imaging

Ten microliter samples were loaded onto a glow-discharge copper grid (200 mesh Cu, Formvar/Carbon 01800-F, Pelco). After absorption for 1 min, excess fluid was drained off by filter paper (Whatman Inc., USA). The grids were stained with 2% uranyl acetate for 1 min and drained off again. The grid was dried under the condition of 30% humidity for 24 hrs. Images were recorded at 30,000X and 50,000X magnification under a transmission electron microscope (TEM, Tecnai G2 Spirit TWIN, FEI Company). The image processing was performed with the EMAN2 software package (Lin, Chang et al., 2013).

### Surface plasmon resonance

All measurements were performed on a BIAcore T200 instrument (GE Healthcare) using a C1 chip (Salanga, Dyer et al., 2014). The chip was clean with two 1 min injections of freshly prepared 0.1 M glycine-NaOH, pH 12 containing 0.3% Triton X-100. The C1 chip was activated with EDC/NHS, blocked with ethanolamine and coated with neutravidin (Invitrogen) (0.2 mg/mL in 20 mM sodium acetate, pH 6.0), and washed with regeneration buffer (0.1 M glycine, 1 M NaCl, 0.1% Tween-20, pH 9.5). A solution of 8 µg/ml biotinylated heparin in running buffer was injected at 5 µl/min for 300 sec. The streptavidin-coated surface without immobilized heparin was used as a reference. All experiments were performed by 300 sec injection and 360 sec dissociation followed by 2.0 M NaCl for 30 sec to regenerate the chip surface. The affinity of the chemokine-heparin interaction was performed by passing six concentrations of chemokines over the chip with 30 µl/ml running buffer (10 mM HEPES, 150 mM NaCl, 3 mM EDTA, 0.05% Tween-20, pH 7.4) at 25 °C. Data were processed with BIAevalution software (GE Healthcare) and analyzed by Scatchard plot to determine the heparin-binding affinity (*K*_*D*_).

### Structure coordinate

The solution structure of mCCL5 was deposited in the Protein Data Bank with the accession code 5YAM and the hCCL5 X-ray trimer structure with the accession code 6AEZ.

## Acknowledgements

The authors are grateful for the NMR facility at National Tsing Hua University and X-ray beamlines of BL13B1 and BL13C1 at the National Synchrotron Radiation Research Center. This work was supported by Taiwan National Science Council (NSC) grants and the Program for Translational Innovation of Biopharmaceutical Development – TSPA, Taiwan.

